# The Forest, the Trees, and the Phylo-diversity Jungle

**DOI:** 10.1101/063461

**Authors:** Florent Mazel, Caroline M. Tucker, Marc W. Cadotte, Silvia B. Carvalho, T. Jonathan Davies, Susanne A. Fritz, Rich Grenyer, Matthew R. Helmus, Arne Ø. Mooers, Sandrine Pavoine, Oliver Purschke, Dan. F. Rosauer, Marten Winter

**Affiliations:** Univ. Grenoble Alpes & CNRS, Laboratoire d’Écologie Alpine, F-38000 Grenoble.; Centre d’Ecologie Fonctionnelle et Evolutive, CNRS, 1919 Route de Mende, 34293 Montpellier Cedex 5, France.; Biological Sciences, University of Toronto-Scarborough, 1265 Military Trail, Scarborough, Ontario, Canada M1C 1A4.; Stake Key Laboratory of Biocontrol, Key Laboratory of Biodiversity Dynamics and Conservation of Guangdong, Higher Education Institutes, College of Ecology and Evolution, Sun Yat-sen University, Xin Gang Xi Lu No. 135, Haizhu District, Guangzhou, PR China; CIBIO/ InBIO, Centro de Investigação em Biodiversidade e Recursos Genéticos da Universidade do Porto, R. Padre Armando Quintas, 4485-661 Vairão, Portugal.; Department of Biology, McGill University, 1205 Avenue du Docteur-Penfield, Montréal, Quebec, Canada H3A 1B1.; African Centre for DNA Barcoding, University of Johannesburg, APK Campus, PO Box 524, Auckland Park 2006, Johannesburg, South Africa; Senckenberg Biodiversity & Climate Research Centre (BiK-F), Senckenberganlage 25, 60325 Frankfurt am Main, Germany.; Institute of Ecology, Evolution and Diversity, Goethe University, Max-von-Laue-Straβe 9, 60438 Frankfurt, Germany; School of Geography and the Environment, University of Oxford, South Parks Road, Oxford, OX1 3QY United Kingdom.; Department of Ecological Sciences - Animal Ecology, Vrije Universiteit, 1081 HV Amsterdam, Netherlands.; Center for Biodiversity, Department of Biology, Temple University, Philadelphia, PA 19122 USA; Department of Biology, Simon Fraser University, 8888 University Drive, Burnaby, BC, Canada V5A 1S6.; Centre of Ecology and Conservation Sciences (UMR 7204 CESCO), Museum National d’Histoire Naturelle, 61 rue Buffon, Paris, France.; Biodiversity Synthesis Research Group, German Centre of Integrative Biodiversity Research, Halle-Jena-Leipzig, Deutscher Platz 5e, DE-04103 Leipzig, Germany; Department of Computer Science, Martin-Luther-University, Halle-Wittenberg, Von-Seckendorff-Platz 1, DE-06120 Halle(Saale), Germany; Geobotany and Botanical Garden, Institute of Biology, Martin Luther University, Halle-Wittenberg, Am Kirchtor 1, DE-06108 Halle (Saale), Germany; Research School of Biology, Australian National University, Acton ACT 2601 Australia

## Abstract

The joint use of phylogenetic trees and ecological data has proven useful for many aspects of ecology. However, there are a multitude of phylo-diversity metrics with complex interdependencies and mathematical redundancies (the so-called ‘jungle’ of metrics). Several recent papers have been trying to ‘map’ this jungle but appear at a first glance to contradict each other. We suggest that these contradictory results are in fact complementary and reflect two approaches to understand diversity metrics: the first focuses on general mathematical properties,the second focuses on assessing metric performance in relation to particular questions.In this manuscript, we discuss the complementarity of the two approaches and in particular how recent papers fit into this categorisation.

## Main text

The joint use of phylogenetic trees and ecological data has proven useful for understanding the assembly of local communities (Webb et al., 2002; Bryant et al., 2008), for exploring large scale diversity patterns (Graham & Fine, 2008), and even for developing target conservation priorities (Isaac et al., 2007). However, the last decades have seen a proliferation of phylo-diversity metrics; we counted more than 70 (Tucker et al. 2016). Winter et al. (2013) refer to the ever-increasing portfolio of phylo-diversity indices as a “jungle”, alluding to both the multitude of metrics and their complex interdependencies and mathematical redundancies. Several groups (e.g. Vellend et al. 2010, Pavoine and Bonsall 2011, Pearse et al. 2014, Tucker et al. 2016) have tried to navigate through this jungle by exploring metric relationships, complementarity and utility. In a recent paper,Miller et al. (2016) contribute an important discipline-specific perspective that further resolves the emerging map.

Metrics can be analysed in two ways: (1) by grouping them based on their underlying properties (e.g. by comparing mathematical formulations); and (2) by assessing context-dependentbehaviour (e.g. by comparing metric performance in relation to particular questions). The first approach requires theoretical and cross-disciplinary studies to summarize the main dimensions along which phylo-diversity metrics vary, while the second provides a field-specific perspective to quantify the ability of a particular metric to test a particular hypothesis. These two approaches have different aims, and their results are not *a priori* expected to be identical.

Miller et al. (2016) carry out this second approach within the discipline of community ecology by testing the ability of 32 phylo-diversity metrics and nine null models in discriminating between two ecological processes: habitat filtering and competition (see e.g. Hardy 2008).

The authors first simulated communities under three main assembly rules: competitive exclusion leading to species being less related than expected by chance, habitat filtering leading to species being more closely related than expected by chance, and neutral assembly. They then tested *a posteriori* which combination of metrics and null models yielded the best statistical performance. Surprisingly, only a fraction of phylo-diversity metrics and null models exhibited the ideal statistical properties of high statistical power coupled with low Type I error rate, leading Miller et al. to conclude that some metrics and null models proposed in the literature should be avoided when asking if filtering and competition play an important role in structuring communities. This is an important finding.

But before theoreticians run off to find new metrics, more multidisciplinary work is in order. One reason there are so many metrics is that they have been pooled across community ecology, macroecology and conservation biology. The questions typically asked by conservationists and macroecologists, for example, differ from those of community ecologists. Different metrics might perform better or worse for different types of problems. One solution would be to explicitly simulate the processes of interest for a given research question (e.g. vicariance or diversification processes in macroecological research), and select the most appropriate metric for the task. The R package presented by Miller *et al.*, as well as others (e.g. Pearse et al. 2015) help facilitate this approach. Of course, applying all possible metrics to the appropriate simulations and null models for a given hypothesis, is complex, time consuming and inefficient. To the extent this is true, this motivates the other approach to navigating the jungle, the unified framework.

We (Tucker et al., 2016) recently took the first approach to metric analysis and classified the 70 phylo-diversity metrics along three broad dimensions: *richness*, *divergence* and *regularity* the sum, mean and variance of phylogenetic distances among species of assemblages, respectively-. Building upon a previous phylo-diversity classification system (Pavoine & Bonsall, 2011), which itself is based on a system for classifying taxonomic and functional diversity metrics (e.g. Ricotta 2007, Villéger et al. 2008), the Tucker et al. (2016) classification offers clear theoretical linkages between phylogenetic and functional approaches in ecology. We then used simulations to corroborate this classification system. Although they conclude differently, we feel that Miller et al. actually provide independent support of this tri-partite classification system. Indeed, the vast majority of metrics used by Miller et al. on their simulated communities group according to this richness-divergence-regularity classification system (see Figure 1 of Miller et al.). And metrics like H_AED_ and E_ED_, which stem from a mathematical combination of richness and regularity dimensions, are expected to sometimes cluster with *richness* (as observed by Miller et al.), and sometimes with *regularity* (see the specific discussion on these hybrid metrics in Tucker et al., 2016).

So, while the synthesis presented in Tucker et al. (2016) takes a broad perspective across fields and across most metrics by providing an objective conceptual classification system, more focussed analyses, such as that by Miller et al., offer a detailed description of metric performance relevant to a given biological question. Both approaches have utility, and importantly, both approaches benefit each other. On one hand, detailed analyses of metric performance offer a valuable test of the broader classification system, using alternative simulations and codes. On the other hand, broad syntheses offer a conceptual framework within which results of more focussed analyses may be interpreted. For example, Miller et al. find that, in some situations, the metrics with the best statistical performances are Rao’s quadratic entropy and IntraMPD. These are hybrid *richness* plus *divergence* metrics. And indeed, the finding that metrics closely aligned with only a single dimension (called ‘anchor’ metrics in Tucker et al. 2016) do not act as the best indicators of community assembly algorithms, provides strong support for the idea that the community assembly processes modelled by Miller et al. actually involve multiple phylo-diversity dimensions. This is a characteristic that has also been recognized in the functional trait literature (see, e.g. Botta-Dukát and Czúcz 2016).

In summary, Miller et al.’s community ecology study offers an excellent complement to broad-scale syntheses such as that provided by Tucker et al. (2016). We call on researchers to continue to hack away at the jungle of phylo-diversity metrics in their own fields, in the hopes that the combination of in-depth understanding of what the metrics are (e.g., how they capture richness, divergence, regularity and combinations thereof), and how they perform under particular ecological and evolutionary processes will allow us a clearer view of our respective fields. Machetes up!

